# TACAN is an essential component of the mechanosensitive ion channel responsible for pain sensing

**DOI:** 10.1101/338673

**Authors:** L. Beaulieu-Laroche, M. Christin, AM Donoghue, F. Agosti, N. Yousefpour, H. Petitjean, A. Davidova, C. Stanton, U. Khan, C. Dietz, E. Faure, T. Fatima, A. MacPherson, A. Ribeiro-Da-Silva, E. Bourinet, R. Blunck, R. Sharif-Naeini

**Affiliations:** Department of Physiology and Cell Information Systems, McGill University.; Department of Physics, Université de Montréal.; Institut de Génomique Fonctionnelle, Université de Montpellier, CNRS, INSERM, Montpellier, France.; Department of Pharmacology and Therapeutics, McGill University.; Department of Pharmacology and Physiology, Université de Montréal.; Department of Brain & Cognitive Sciences and McGovern Institute for Brain Research, Massachusetts Institute of Technology.; Alan Edwards Center for Research on Pain.

## Abstract

Mechanotransduction, the conversion of mechanical stimuli into electrical signals, is a fundamental process underlying several physiological functions such as touch and pain sensing, hearing and proprioception. This process is carried out by specialized mechanosensitive ion channels whose identities have been discovered for most functions except pain sensing. Here we report the identification of TACAN (Tmem120A), an essential subunit of the mechanosensitive ion channel responsible for sensing mechanical pain. TACAN is expressed in a subset of nociceptors, and its heterologous expression increases mechanically-evoked currents in cell lines. Purification and reconstitution of TACAN in synthetic lipids generates a functional ion channel. Finally, knocking down TACAN decreases the mechanosensitivity of nociceptors and reduces behavioral responses to mechanical but not to thermal pain stimuli, without affecting the sensitivity to touch stimuli. We propose that TACAN is a pore-forming subunit of the mechanosensitive ion channel responsible for sensing mechanical pain.

## Introduction

The sensations of touch and pain are essential to our interaction with our environment. They rely on a cellular phenomenon called mechanotransduction, during which mechanical forces are converted into electrical signals. This vital process underlies several physiological functions such as hearing, touch, pain, and proprioception (Boyer et al., 2011; Chalfie, 2009; Li et al., 2012; Woo et al., 2015), as well as the myogenic tone of resistance arteries (Sharif-Naeini et al., 2009), fluid homeostasis and the sensation of thirst (Bourque, 2008; Zaelzer et al., 2015), and flow sensing in kidney tubules (Peyronnet et al., 2012). Many of these functions require mechanotransduction at the micro-to millisecond scale, suggesting that the transducers are mechanosensitive ion channels (MSCs). This is confirmed by the recent discovery of genes responsible for our senses or touch (Ranade et al., 2014b), hearing (Pan et al., 2013), and proprioception (Boyer et al., 2011; Woo et al., 2015). Yet the genes responsible for our most unwanted experience of mechanotransduction, our sense of pain, remain elusive. Here, we report the identification of an essential component of the mechanosensitive ion channel in pain-sensing neurons. We find that TACAN is a pore-forming subunit that mediates mechanotransduction in nociceptors and is necessary for sensing mechanical pain.

## Results

### TACAN is a candidate mechanosensitive ion channel found in nociceptors

A previous proteomic screen of membrane proteins involved in mechanotransduction in smooth muscle cells generated a list of over 70 candidates (Sharif-Naeini et al., 2009), including some transmembrane proteins of unknown function. One of these candidates, which we named TACAN (for “movement” in Farsi), also known as TMEM120A, NET29 or TMPIT, is a highly-conserved protein from humans to zebrafish (Figures 1A-1B and S1). The predicted membrane topology (Constrained Consensus TOPology (CCTOP)) suggests TACAN has 6 transmembrane domains with internal amino- and carboxy- termini (Figure 1C). Tissue expression analysis reveals that TACAN is expressed in several tissues (Figure 1D), with a higher expression in heart, kidney, colon, and sensory neurons of the dorsal root ganglia (DRGs). Given the role of DRG neurons in our senses of touch and pain, we examined whether the expression of TACAN in this tissue was restricted to a specific subset of neurons. *In situ* hybridization experiments (Figure 2A) demonstrated the presence of TACAN mRNA in a broad set of DRG neurons. Analysis of the cross-sectional area distribution, however, revealed that TACAN expression is biased toward small to medium diameter neurons (Figure 2B), suggesting its expression could be localized in nociceptors. To examine the subset of sensory neurons expressing TACAN, we generated a rabbit anti-TACAN antibody (Figure S2). Our data indicate that TACAN is highly expressed in non-peptidergic nociceptors, based on colocalization with markers such as P2×3, GINIP and IB4 (Figures 2C and 2D). Colocalization was low with markers of myelinated neurons (NF200), peptidergic nociceptors (CGRP), or proprioceptors (parvalbumin). Tyrosine hydroxylase, a marker of low-threshold C-fibers, was found in about a third of TACAN-positive sensory neurons. A previous study reported the expression of TACAN on the nuclear membrane of adipocytes (Batrakou et al., 2015). We verified the plasma membrane-localization of HA-tagged TACAN in CHO cells and the sensory neuron-derived F11 cells. Our immunofluorescence and membrane biotinylation data indicate TACAN is present at the plasma membrane (Figure S2).

**Figure 1.**
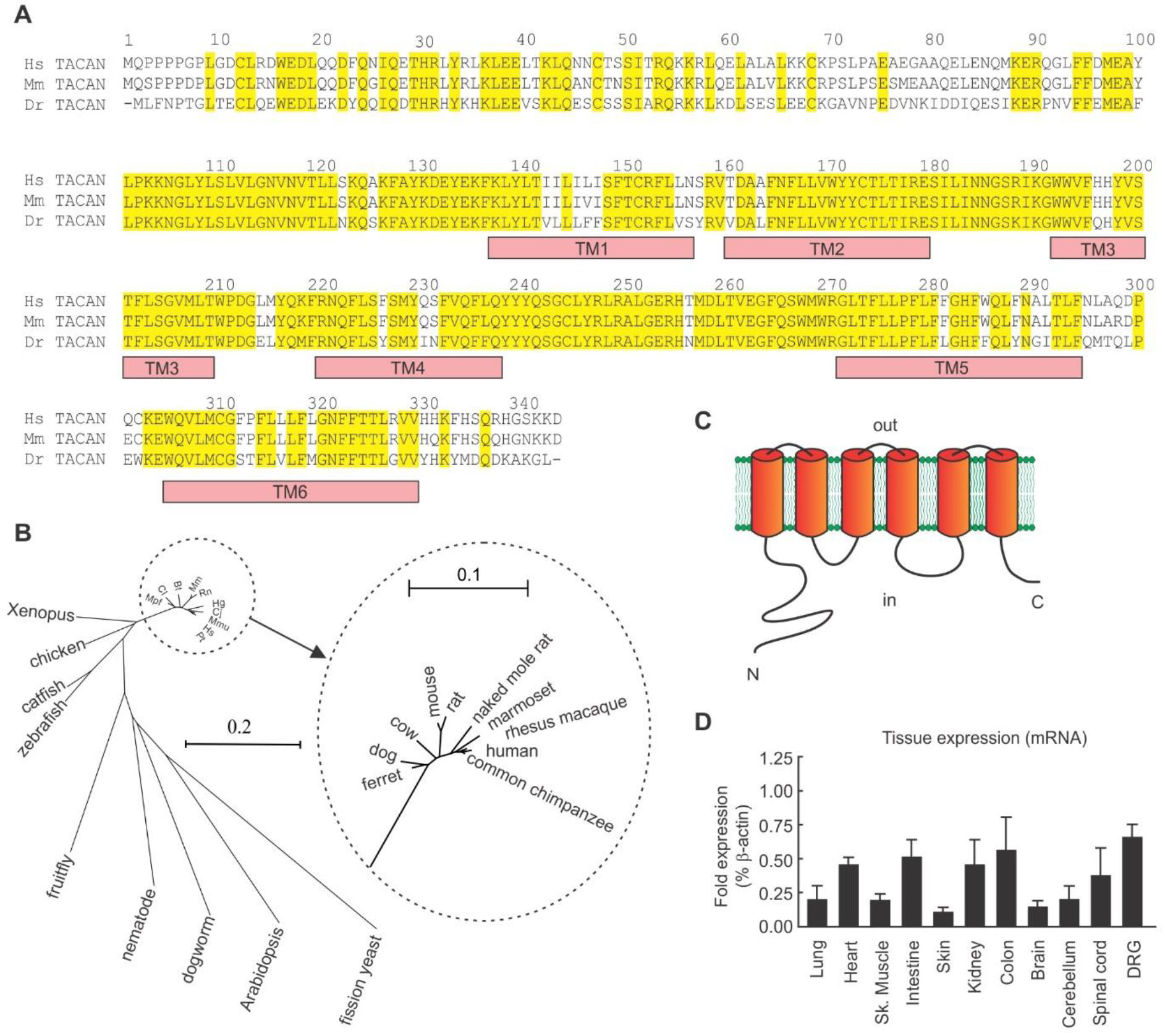
Analysis of TACAN amino acid sequence, phylogenic tree and predicted membrane topology. (A) Comparison of the amino acid sequences for TACAN from human (Hs), mouse (Mm) and zebrafish (Dr). Amino acid residues are numbered from the first methionine of Hs TACAN. Conserved residues are highlighted in yellow, predicted transmembrane (TM1-TM6) segments are indicated by the orange boxes, as predicted by the CCTOP server. (B) Unrooted phylogenic tree showing sequence relationship of TACAN in different species. Alignments were generated by Clustal-Omega and phylogenic tree generated using Unrooted (http://pbil.univ-lyon1.fr/software/unrooted.html). Scale: number of substitutions per position. (C) Predicted membrane topology of TACAN.

**Figure 2.**
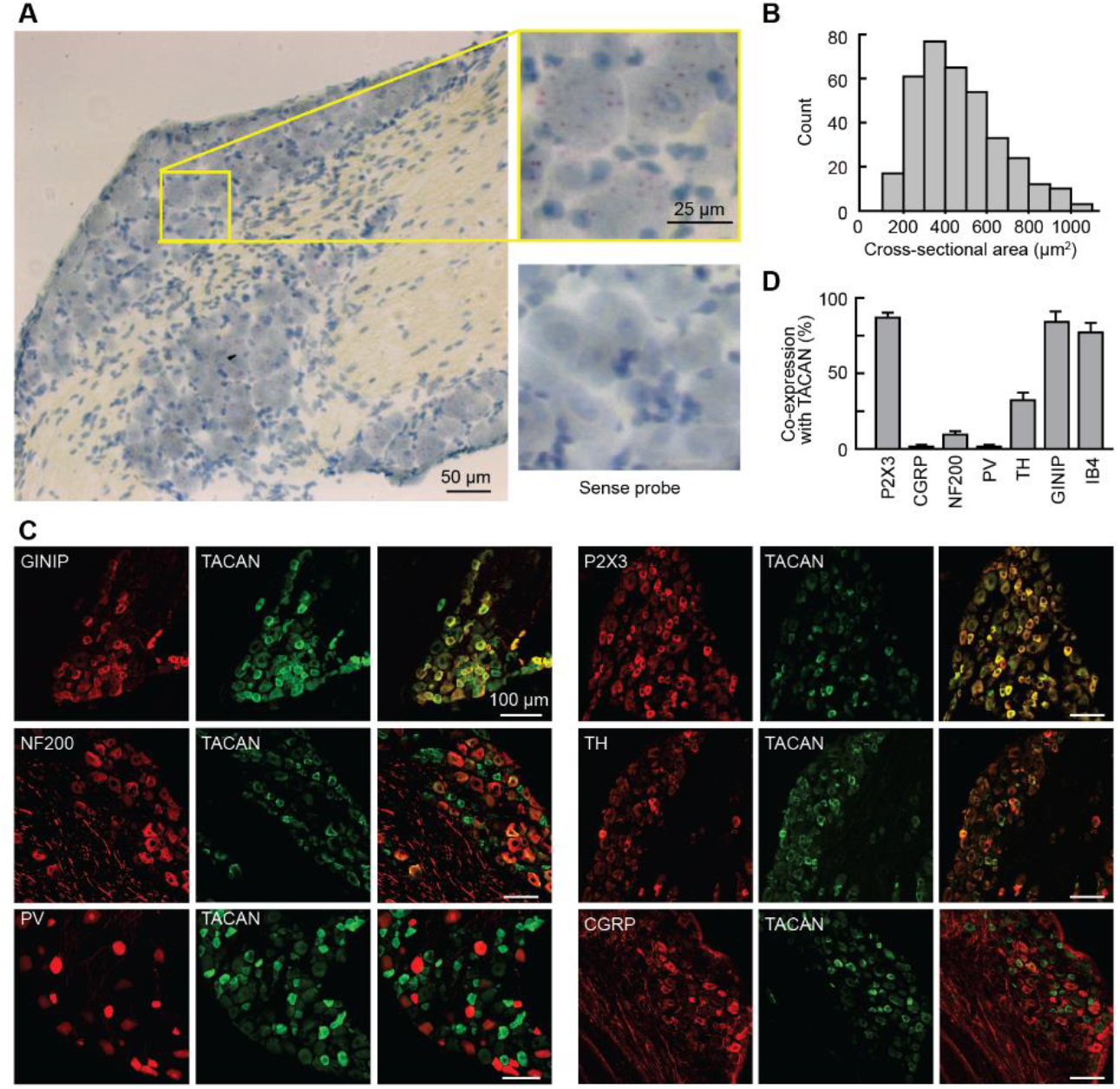
TACAN is expressed in nociceptors. (A) *In situ* hybridization examining the presence of TACAN in mouse DRG sections. Top right panel is a magnification of the box in the left panel, with the TACAN probe. Bottom right panel is a magnification of a DRG section probed with a sense (control) probe. (B) Cross-sectional area distribution of TACAN mRNA-positive neurons. (C) Double immunostaining to determine the colocalization of TACAN with markers of myelinated afferents (Neurofilament 200; NF200), peptidergic (CGRP) and non-peptidergic (P2×3) nociceptors, C-type low threshold mechanoreceptors (C-LTMR, Tyrosine hydroxylase TH), and GINIP (C-LTMRs and non-peptidergic nociceptors). Colocalization with proprioceptors was done by staining DRG sections of *Pvalb:cre; tdTom* mice with the TACAN antibody. (D) Percentage (±s.e.m.) of marker-immunoreactive neurons that co-express TACAN.

### TACAN expression increases mechanically-evoked currents in heterologous systems

A criterion for channel mechanosensitivity is that its expression in heterologous systems increases mechanically-evoked currents. In CHO cells stably transfected with TACAN, we used cell-attached recordings to determine whether this candidate increases mechanically-evoked currents (Figures 3A and 3B). To activate MSCs in the membrane patch recorded, we applied brief pulses of negative pressure through the recording electrode. In MOCK-transfected CHO cells, we observed the presence of endogenous MSCs (Figure 3A, left panel). Stable expression of TACAN in CHO cells significantly increased mechanically-evoked currents when compared to MOCK-transfected cells (Figures 3A and 3B). Although the activation threshold of the channels did not change (−33.9 ± 2.7 mm Hg, n=31, vs. −35.1 ± 3.1 mm Hg, n=55, in MOCK- and TACAN-expressing cells, respectively), the percentage of active patches was significantly higher (53% vs 91% in MOCK-vs. TACAN-expressing cells; p<0.001; Fisher exact test), suggesting a higher density of MSC.

**Figure 3.**
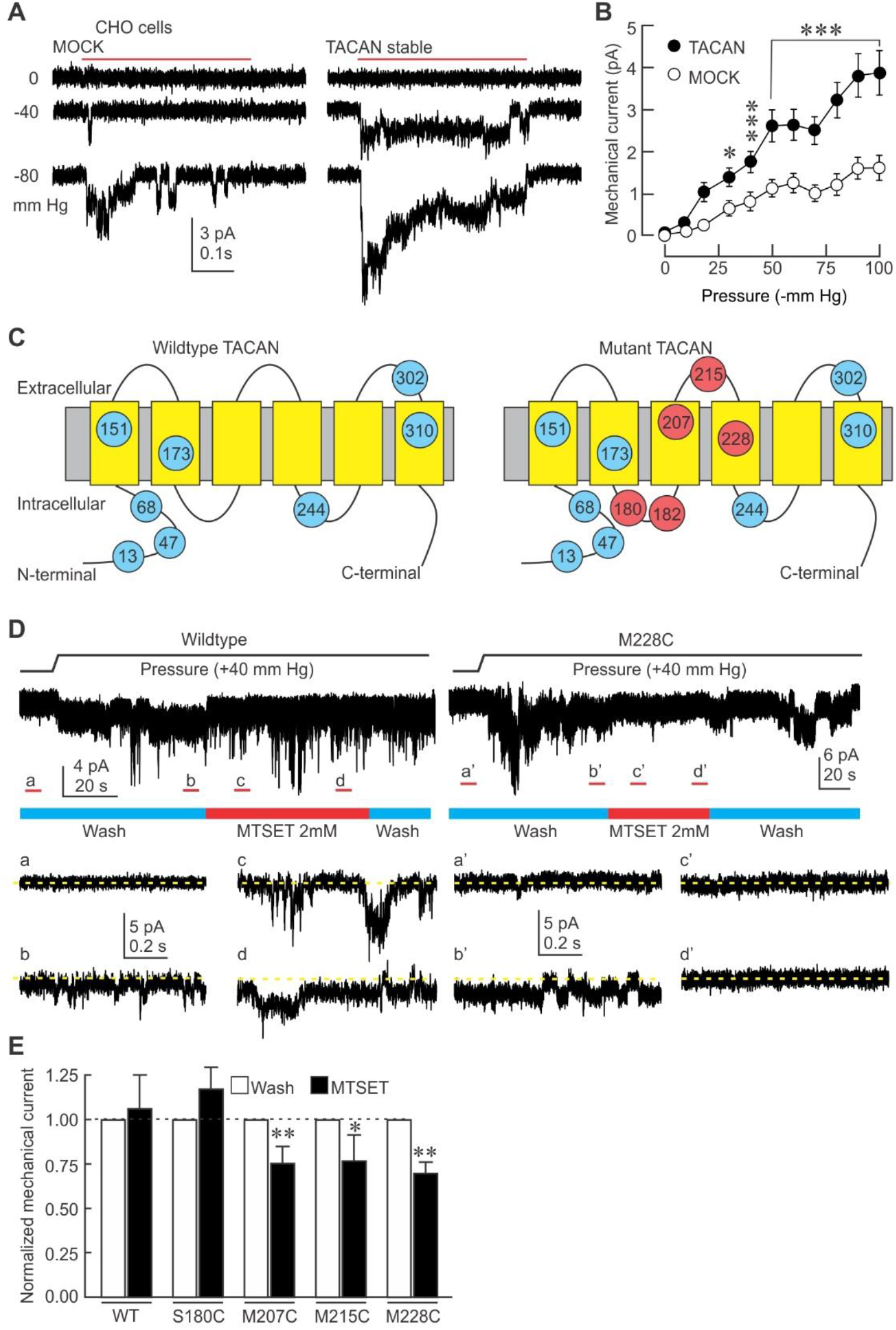
TACAN is an essential part of a mechanosensitive ion channel in heterologous systems. (A) Representative cell-attached current traces elicited by mechanical stimuli (red bar) in CHO cells stably expressing TACAN or MOCK. (B) Mean (±s.e.m.) mechanically-evoked currents in stable CHO cells expressing MOCK (n=58) or TACAN (n=61). *, *** =p<0.05; and 0.001, respectively. Asterisks indicate significant difference from MOCK group (Two-way ANOVA with Tukey post-hoc test). (C) Predicted membrane topology of TACAN with its endogenous cysteine residues (blue, amino acid number indicated in circles), and the residues that were individually substituted (red). (D) Left: representative current trace of an outside-out patch excised from a HEK293 cells transfected with WT TACAN and stimulated with positive pressure through the recording electrode. Insets represent the non-stimulated baseline period (a), stimulated channels (b) in the patch superfused with control solution (wash, blue bar), in stimulated channels exposed to MTSET (2mM, red bar) early (c) and late (d) in the exposure. Dotted yellow line indicates zero current level. Right: same protocol with HEK293 cells transfected with M228C TACAN. (E) Mean (±s.e.m.) normalized mechanically-evoked current in WT and mutant HEK293 cells. (Paired t test, * = p<0.05; ** p<0.01).

To determine if this increase in cell mechanosensitivity occurs in other cell types, we examined the effect of TACAN expression on mechanically-evoked currents in COS-7 and HEK293 cells (HEK293/17) (Figures S3A-S3D), which are commonly used to study MSCs (Anderson et al., 2018; Coste et al., 2010; Li Fraine et al., 2017). In MOCK-transfected COS7 cells, we observed the presence of endogenous MSCs (Figure S3A, left panel) with a low activation threshold, often as low as −5 mm Hg (mean activation threshold: −21.3 ± 1.6 mm Hg, n=97), consistent with previous reports (Sharif-Naeini et al., 2009). Most recordings (93%, 90 out of 97 patches) had active MSCs in them. After transfecting these cells with TACAN, we observed once again a significant increase (p<0.05; Two-Way ANOVA with Tukey post-hoc test) in mechanically-evoked currents. In HEK293 cells, endogenous MSCs were present in most patches (84%, 66 of 79) and activated at a threshold of −42.0 ± 2.6 mm Hg (n=66) to reach a plateau of activation (Figures S3C and S3D). However, when these cells were transfected with TACAN, an additional increase in mechanosensitive current became evident at thresholds negative to - 80 mm Hg (Figure S3D), suggesting the recruitment of another MSC. This increase in mechanically-evoked current did not affect the channel activation threshold (−37.1 + 3.2 mm Hg, n=47) or the percentage of active patches (82%, 47 of 57 patches). This elevated activation threshold is similar to that observed in cultured primary nociceptors (Cho et al., 2002), also known as high-threshold mechano-nociceptors, highlighting another similarity between TACAN and the endogenous channel found in nociceptors. In stably transfected CHO cells, we determined the single-channel conductance of the MSC induced by TACAN proteins. Our results demonstrate that MSCs in TACAN-expressing CHO cells are characterized by a linear current-voltage (I–V) relationship with a reversal potential of 6.14 ± 2.40 mV (Figure S3E and S3F, n=11). This is consistent with this MSC mediating a non-selective cationic conductance. These I–V relationships resulted in slope-conductance values in these recording conditions of 11.5 ± 0.5 pS for TACAN cells (n= 11). Single-channel recordings with only CaCl_2_ as the charge carrier demonstrated this channel is permeable to calcium ions as well (Figure S3G). We further demonstrated through whole-cell recordings that, in the absence of mechanical stimuli, TACAN expression did not impact the voltage-gated conductances (Figure S3H). Although TACAN expression led to an increase in mechanically-evoked currents in all three cell lines, the profile of the current was different. This could be due to TACAN’s association with different accessory subunits, or to the different visco-elastic properties of the different membranes.

Endogenous MSCs are under tonic inhibition by the F-actin cytoskeleton, and destabilizing agents such as cytochalasin (Cyto-D) can increase mechanically-evoked currents (Sharif-Naeini et al., 2009). We examined the possibility that the enhanced mechanical responses in our TACAN stable cell line are due to TACAN disrupting the cytoskeletal inhibition of the endogenous MSC rather than TACAN being a part of the mechanosensitive channel complex. If this was the case, we would expect Cyto-D to significantly increase mechanically-evoked currents in control cells, but have little effect in TACAN-expressing cells. In control CHO cells, treatment with Cyto-D (1 μM) significantly increased mechanically-evoked currents (compared to vehicle) to a level similar to that of vehicle-treated cells of the TACAN stable cell line (Figure S3I). However, Cyto-D treatment of TACAN-expressing cells also led to a significant increase in mechanically-evoked currents, indicating that cytoskeletal inhibition of MSCs is still present in these cells and is not responsible for their increased mechanosensitivity (Figure S3I).

Our results indicate that TACAN expression increases cellular mechanosensitivity, but it remains unclear whether TACAN is an essential part of a novel MSC or potentiates endogenous MSCs. To examine the former possibility, we used the substituted-cysteine accessibility method (SCAM) to introduce single cysteine mutations in TACAN at sites predicted to be near the extracellular side and near a potential ion-conducting pore (MEMSAT-SVM prediction (http://bioinf.cs.ucl.ac.uk/psipred)) (Figure 3C). The sulfhydryl group of these cysteines can then be targeted by the thiol-reactive reagent [2-(trimethylammonium)ethyl] methanethiosulfonate bromide (MTSET, 2mM). We hypothesized that if TACAN is part of a novel MSC, the mechanically-evoked currents in TACAN-transfected HEK293 cells would be significantly reduced by the application of MTSET. In HEK293 cells transfected with WT TACAN, which possesses several endogenous cysteine residues (see blue circles in Figure 3C), including one in the extracellular loop between the fifth and sixth transmembrane domains (S302), application of MTSET to mechanically-stimulated outside-out patches had no effect on the activity of MSCs (Figure 3D, upper panel). However, when HEK293 cells were transfected with TACAN bearing cysteine residues in the third or fourth transmembrane domains, or in the extracellular loop connecting them, application of MTSET caused a significant reduction of the mechanically-evoked currents (Figures 3D and 3E). These mutations did not affect the membrane targeting of TACAN, assessed by cell-attached single-channel recordings, or membrane biotinylation experiments (data not shown). As expected, when the cysteine residue is placed in the intracellular loop between the second and third transmembrane domains, MTSET did not reduce mechanically-evoked currents. These results indicate that TACAN forms the ion-conducting part of a MSC responsible for mechanically-evoked currents in these cells.

### TACAN forms an ion-conducting channel when reconstituted in artificial bilayers

It was recently suggested that some candidate MSCs may act as permissive factors that promote the expression or membrane targeting of the endogenous channel (Anderson et al., 2018; Dubin et al., 2017). Alternatively, they may need additional subunits to form a functional ion channel. To determine whether the expression of TACAN proteins alone was sufficient to generate an ion-conducting pore, we reconstituted isolated TACAN into lipid bilayers formed of synthetic lipids. After purification (Methods) and reconstitution in planar lipid bilayers, we measured the current across the membrane when voltage steps were imposed (Figure 4). Following Laplace’s Law (Sachs, 2010), membranes with a high radius of curvature, such as those in planar bilayers, should hold the MSCs in the open configuration and allow them to flux ions. The addition of TACAN-containing proteoliposomes to the planar bilayer produced macroscopic currents (Figure 4A), indicating that TACAN can by itself form an ion-conducting pore. Furthermore, the TACAN protein-elicited currents displayed Ohmic behavior (Figure 4B), similar to the one observed in the stable CHO cells. Finally, the TACAN channel was blocked by known blockers of MSCs such as gadolinium (30 μM), which reduced the current down to 21% and 36% of baseline at +80mV and −80mV, respectively. The peptide GsMTx4 (5 μM; Figures 4C and 4D), a MSC blocker purified from the venom of a Chilean rose tarantula, *Grammostola spatulata* (Bae et al., 2011; Suchyna et al., 2000), reduced the current down to <1% of baseline. These blockers were used at similar concentrations known to inhibit the endogenous MSCs (Cho et al., 2006; Drew et al., 2002; Sharif-Naeini et al., 2009; Yang and Sachs, 1989). Our results indicate that TACAN forms an ion-conducting pore that is sensitive to known blockers of MSCs.

**Figure 4.**
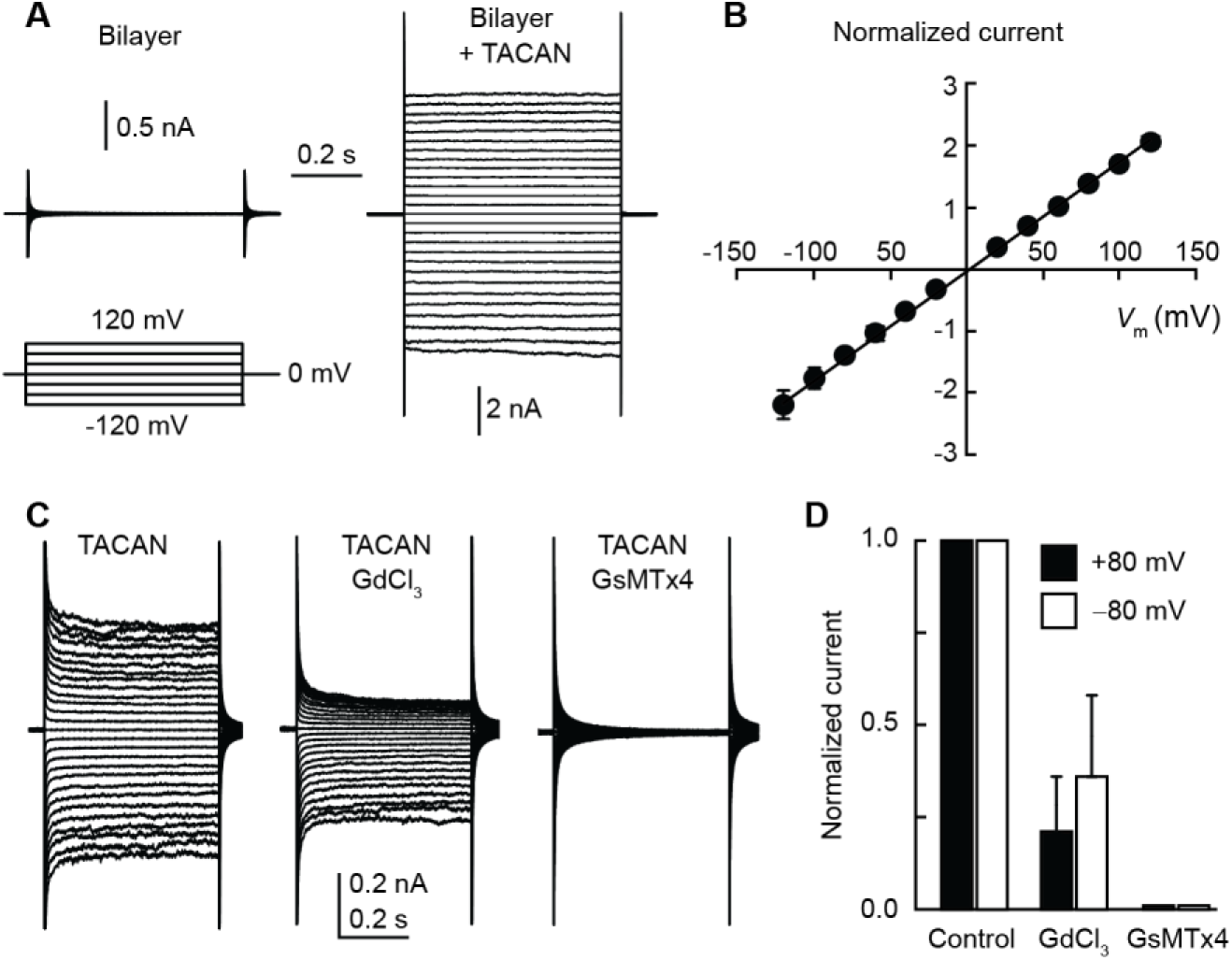
Macroscopic currents elicited from isolated TACAN channels in planar lipid bilayer. (A) A planar lipid bilayer showing no conductance prior to incorporation (left panel) develops macroscopic currents (right panel) after insertion of purified and reconstituted TACAN channels. Traces show currents in response to voltage pulses from a resting value of 0 mV to voltages between −120 and +120 mV in intervals of 10 mV. (B) Mean (±s.d.; n=7) current normalized to the current value at +80 mV. The I-V characteristic is consistent with a non-selective cation channel as observed in whole-cell and cell-attached recordings. (C) The conductance is sensitive to gadolinium (GdCl_3_; 30 μM) and GsMTx4 (5 μM). (D) Bilayer current (normalized to pre-blocker current) after addition of GdCl_3_ and GsMTx4.

### TACAN is essential to the intrinsic mechanosensitivity of nociceptors

We next asked whether TACAN underlies the mechanosensitivity of nociceptors. Sensory DRG neurons were cultured from reporter mice in which nociceptors were visually-identifiable through the expression of tdTomato. Neurons were transfected with Alexa488-labeled siRNA molecules targeting TACAN or a non-targeting (NT) control (Figure 5A). The knockdown was validated using a neuroblastoma (N2A) cell line (Coste et al., 2010) (Figure S5). Our data demonstrate that reducing the expression level of TACAN led to a significant reduction in mechanically-evoked currents in nociceptors (Figures 5B-5D). This decrease was accompanied by a significant reduction in the number of patches with active MSCs (44/54 (81%) in NT-siRNA compared to 40/67 (59%) in TACAN-siRNA-transfected nociceptors; Fisher Exact probability test, p=0.008). This led us to examine whether GsMTx4, which inhibits purified TACAN-elicited currents in synthetic lipids, could also inhibit mechanically-evoked currents in nociceptors. Our data demonstrate that, in outside-out recordings from DRG neurons, superfusion of GsMTx4, used at doses that blocked TACAN-elicited currents in the bilayer (5 μM), led to a significant and reversible reduction of mechanically-evoked currents (Figures 5E and 5F).

**Figure 5.**
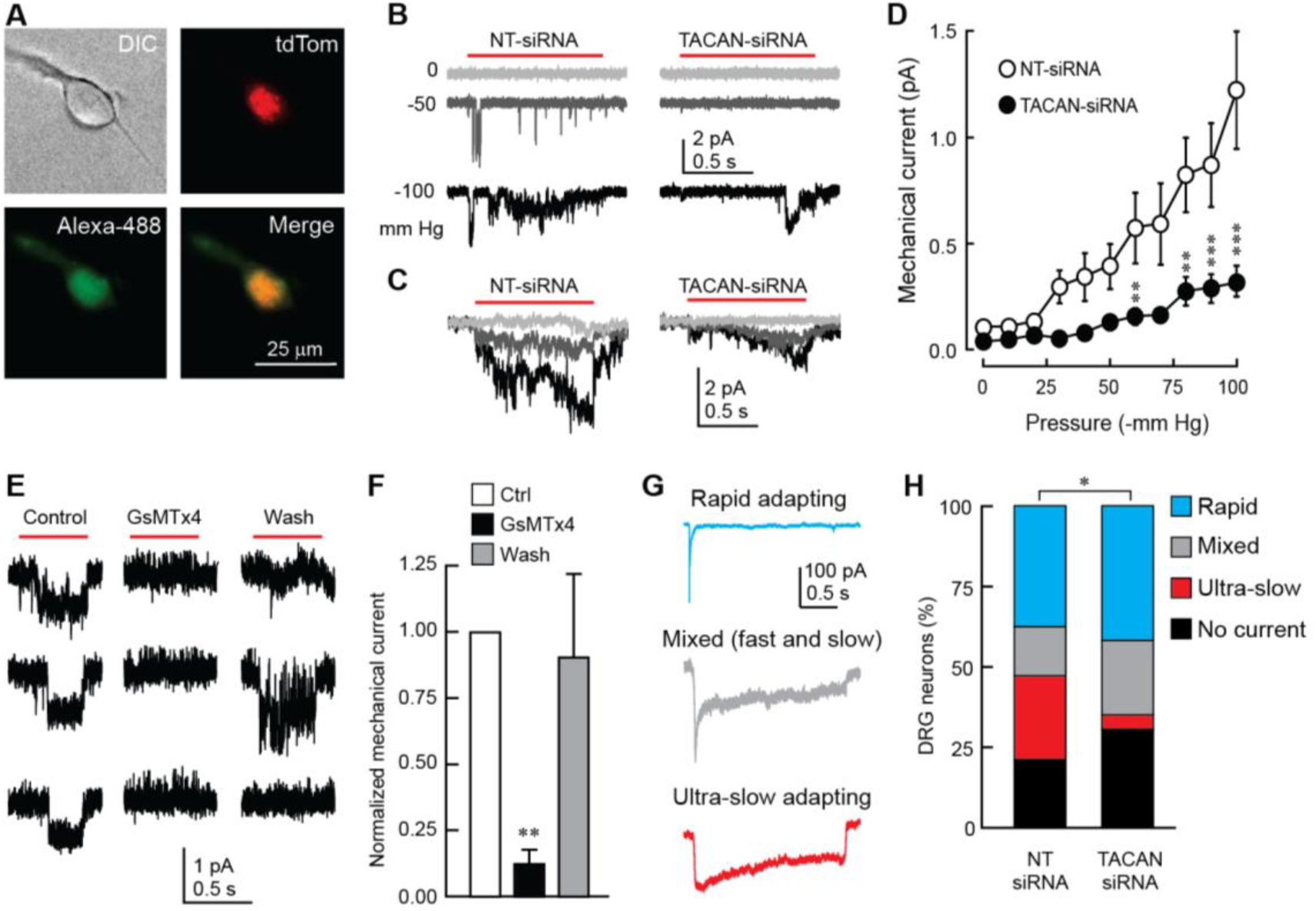
TACAN is essential to the mechanosensitivity of nociceptors. (A) Representative image of a cultured nociceptor from a *Trpv1-tdTom* mouse transfected with Alexa 488-conjugated siRNA molecules targeting TACAN. (B) Representative mechanically-evoked cell-attached currents in cultured nociceptors 72 hours after transfection of control- (NT-siRNA) or TACAN-targeting siRNA molecules. (C) Average current traces of mechanically-evoked responses in nociceptors transfected with NT- or TACAN-siRNA (n=77 and 66 cells, respectively). (D) Mean (±s.e.m.) mechanically-evoked currents in nociceptors transfected with NT- or TACAN-targeting siRNA. **, *** =p<0.01; and 0.001, respectively. Asterisks indicate significant difference from the NT-siRNA group (Two-way ANOVA with Tukey post-hoc test). (E) Representative mechanically-evoked currents obtained from DRG neurons in the outside-out configuration in response to a positive pressure pulse from the recording pipette (red bar) before (Control), during (GsMTx4) and after (Wash) the application of 5 μM GsMTx4. The three rows represent repetitive trials in each condition. (F) Mean (±s.e.m.) mechanically-evoked currents extracted from the approach depicted in E (n=6). (G) Representative types of mechanically-evoked whole-cell currents from cultured DRG neurons stimulated by an indenting stimulus from a blunt probe. (H) Percentages of neurons displaying the response types shown in G (n = 19 and 26 neurons in NT-and TACAN-siRNA conditions, respectively. *p<0.05; Contingency test with Fisher’s Exact Probability test).

Mechanically-evoked whole-cell currents in nociceptors display a unique non-adapting or ultra-slow adapting kinetic that distinguishes them from the rapid-adapting currents recorded in large-diameter neurons (Hao and Delmas, 2010). The latter has been associated with expression of Piezo2 (Ranade et al., 2014b; Woo et al., 2015). To determine whether TACAN is responsible for the ultra-slow adapting current in nociceptors, we recorded whole-cell mechanically-evoked currents from DRG neurons obtained from wild-type mice, transfected with the same set of Alexa488-labeled siRNA molecules. We defined three types of whole-cell currents based on their inactivation kinetics (Figure 5G) (Francois et al., 2015). Compared to NT-siRNA transfected neurons (n=19), treatment of DRG neurons with TACAN-siRNA (n=26) caused a significant (p<0.05; Fisher’s Exact Probability test) decrease in the proportion of neurons displaying the ultra-slow adapting current (Figure 5H). We observed an increase, yet non-significant, in the proportion of neurons displaying no or mixed currents in the TACAN siRNA-treated group, as predicted if loss of TACAN converts ultra-slow adapting neurons into mixed or non-responders.

### TACAN expression is essential for the detection of painful mechanical stimuli *in vivo*

We next asked whether TACAN is necessary for the detection of painful mechanical stimuli. We hypothesized that reducing TACAN expression would impair responses to high-intensity mechanical stimuli but not to low-intensity stimuli. Furthermore, the response to non-mechanical nociceptive stimuli, such as heat, should be unaffected. We injected AAV2/6 viral particles expressing a shRNA targeting TACAN or NT control directly in the left sciatic nerve bundle. The knockdown of TACAN by the virus was validated in F11 cells (Figure S5). Reducing the expression of TACAN in sensory neurons significantly decreased the mouse withdrawal responses to high intensity von Frey stimuli (Figure 6A). Given that the withdrawal response can be the result of spinal reflexes, we also examined the mouse behavior for evidence of pain-related responses, such as paw flicking or licking, as previously described (Ducourneau et al., 2014). When the intensity of the mechanical stimulus rose above the withdrawal threshold, mice began to display nociceptive-like behaviors (Figure 6B). Interestingly, reducing the expression level of TACAN led to a significant decrease in these nociceptive behaviors. Mice injected with the non-targeting shRNA virus showed no change in their sensitivity to mechanical stimuli, both below and above the withdrawal threshold. The loss of nociceptive behavior in TACAN-knockdown animals was specific to mechanical stimuli, as the same mice continued to exhibit normal nociceptive responses to painful thermal stimuli (Hargreaves’ test, Figure 6C). Finally, reflexive paw withdrawal from a pinprick stimuli, a model of intense mechanical pain stimuli resulting from activation of peptidergic A-*δ* nociceptors (Arcourt et al., 2017), was not different in the two groups (Figure 6D). Taken together, these data indicate that TACAN is required for mice to detect painful mechanical stimuli.

**Figure 6.**
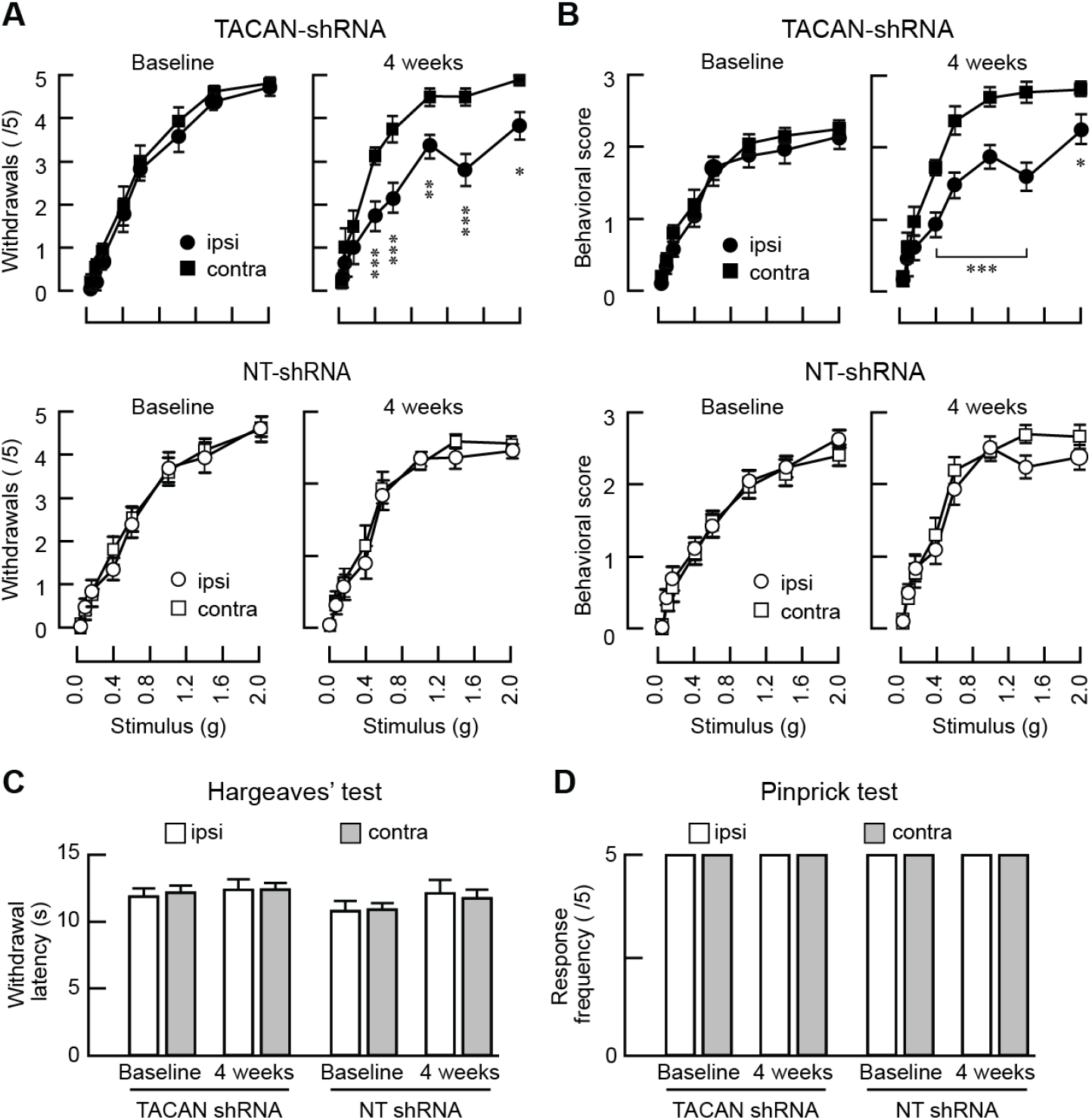
TACAN is essential for the detection of painful mechanical stimuli in vivo. (A) Mean (±s.e.m.) number of mechanically-elicited paw withdrawals in response to von Frey filaments application in mice with sciatic nerve injection of AAVs expressing TACAN-shRNA (top panels, n=13 mice) or NT-shRNA (bottom panels, n=12). Measurements were done before the injection (left) and 4 weeks post-virus injection (right). *, **, *** = p<0.05; 0.01; 0.001, respectively. Asterisks indicate significant difference between ipsilateral and contralateral paws (Two-way RM ANOVA with Bonferroni post-hoc test). (B) Mean (±s.e.m.) nociceptive behavioral scores in response to von Frey filaments application in mice injected with TACAN-shRNA (top panels, n=13) or NT-shRNA (bottom panels, n=12). Measurements were done before the injection (left) and 4 weeks post-virus injection (right). *, *** = p<0.05; 0.001, respectively. Asterisks indicate significant difference between ipsilateral and contralateral paws (Two-way RM ANOVA with Bonferroni post-hoc). (C) Mean (±s.e.m.) withdrawal latencies to radiant heat application (Hargreaves’ test) in mice injected with either the NT-shRNA (n=12) or TACAN shRNA (n=13). No significant changes are observed (ipsilateral vs contralateral, Paired t test). (D) Mean (±s.e.m.) number of withdrawals (out of 5 applications) after mechanical stimuli induced by a pinprick. Neither groups showed a modification of their response to a noxious mechanical stimulus (ipsilateral vs contralateral, Paired t test).

## Discussion

We identified an essential component of a novel mechanosensitive ion channel. TACAN forms a functional channel in lipid bilayers and increases mechanosensitivity in various cell types. We demonstrate that TACAN mediates mechanically-evoked currents in nociceptors and is necessary for pain sensing *in vivo*. A different MSC, termed Piezo2, plays a crucial role in sensing touch but not mechanical pain (Ranade et al., 2014b). Moreover, Piezo2 displays rapid adapting kinetics and a higher conductance than the one identified in nociceptors (Cho et al., 2006; Cho et al., 2002; Coste et al., 2010). In contrast, TACAN displays slow-adapting kinetics, lower conductance, and when its expression is reduced, leaves responses to touch intact but impairs those to mechanical pain. Our results thus indicate that TACAN and Piezo2 have complementary roles in somatosensation.

Interestingly, withdrawal reflexes to an intense painful mechanical stimulus, the pinprick test, were not affected by knocking down TACAN. While this can be explained by a recent report indicating pinprick responses result from the activation of peptidergic A*δ* fibers that express the NPY2 receptor as well as CGRP (Arcourt et al., 2017), it also indicates that peptidergic nociceptors may have different mechanosensitive channels. This is supported by an earlier report indicating the presence of a ~23 pS MSC found in small diameter nociceptors (Cho et al., 2006), although that channel had a low threshold for activation.

TACAN defines a novel class of ion channels that appears specialized to transduce mechanical forces into electrical signals. Much like the Piezo family of MSC, which was found to be involved in touch (Ranade et al., 2014b), proprioception (Woo et al., 2015), lung inflation (Nonomura et al., 2017), vascular development (Ranade et al., 2014a), and red blood cell volume regulation (Cahalan et al., 2015; Ma et al., 2018), TACAN is expressed in many tissues and likely to play important roles in several mechanotransduction processes. TACAN also has a homolog in mammalian cells, TMEM120B, but its heterologous expression did not increase mechanically-evoked currents (Figure S6). Our bilayer reconstitution experiments indicate that TACAN can form an ion channel by itself, but TMEM120B may act as an accessory subunit to TACAN.

Activation of nociceptors is central to the experience of pain, and several chronic pain conditions are caused by the sensitization of nociceptors to mechanical stimuli, including osteoarthritis and rheumatoid arthritis pain (He et al., 2017; McQueen et al., 1991; Neogi et al., 2016; Okun et al., 2012; Schaible, 2014). Intervening at the level of primary afferent nociceptors is a promising approach to developing therapeutic treatments. Yet pain management in many of these chronic inflammatory conditions is currently inadequate in part because the identity of the MSC had remained elusive. With the discovery of TACAN, we have identified a valuable therapeutic target in the treatment of chronic inflammatory pain.

## Acknowledgments

This work was supported by operating grants from the Canadian Institutes of Health Research, pilot grant from the Groupe d’Etude des Protéines Membranaires (GEPROM), and salary award from the Fonds de Recherche du Quebec-Santé (RSN). EB was supported by grants from Fondation pour la Recherche Médicale (équipe FRM 2015) and the Agence Nationale pour la Recherche (ANR15-CE16-0012-01, LABEX ICST). ARS was supported by an operating grant of the Canadian Institutes of Health Research. M.C was supported by a fellowship from the Louise and Alan Edwards Foundation. LBL was supported by an NSERC USRA.

